# Experience shapes the transformation of olfactory representations along the cortico-hippocampal pathway

**DOI:** 10.1101/2024.09.30.615807

**Authors:** Eleonore Schiltz, Martijn Broux, Cagatay Aydin, Pedro Goncalves, Sebastian Haesler

## Abstract

Perception relies on the neural representation of sensory stimuli. Primary sensory cortical representations have been extensively studied, but how sensory information propagates to memory-related multisensory areas has not been well described. We studied this question in the olfactory cortico-hippocampal pathway in mice. We recorded single units in the anterior olfactory nucleus (AON), the anterior piriform cortex (aPCx), lateral entorhinal cortex (LEC), the hippocampal CA1 subfield, and the subiculum (SUB) while animals performed a non-associative learning paradigm involving novel and familiar stimuli. In the AON, neurons were broadly tuned to different chemicals, and their responses were strongly modulated by experience. From the AON to hippocampal structures, the selectivity of neurons for specific odorants increased, concurrent with the development of population-level odor representations, which became independent of novelty and familiarity. While both stimulus identity and experience were thus reflected in all regions, their neural representations progressively separated. Our findings provide a potential mechanism for how sensory representations are transformed to support stimulus identification and implicit memories.

## INTRODUCTION

Sensory systems form a neural representation of the outside world inside the brain by converting physical stimuli into sequences of action potentials. These neural representations enable the detection, identification, and discrimination of stimuli in the environment. The representation of stimuli in primary auditory, visual, somatosensory, and olfactory cortices has been studied extensively (deCharms and Zador, 2000; Kriegeskorte and Wei, 2021), but how stimulus representations transform from primary sensory to multisensory areas is not well described. Moreover, cortical representations are not static, but they undergo changes over shorter (>seconds; Homann et al., 2022; Ulanovsky et al., 2003) and longer (>days; Frenkel et al., 2006; Schoonover et al., 2021; Woloszyn and Sheinberg, 2012) timescales. Nonetheless, non-instrumental stimuli can be remembered in large number (Standing, 1973) and with remarkable detail (Brady et al., 2008). While non-associative learning can thus create vast, implicit memory of stimuli in the environment, it is unclear how stimulus representations reflect this memory.

Here, we characterized the transformation of neural representations from cortical primary sensory to multisensory areas in the lateral entorhinal cortex and hippocampus (LEC-Hpc) and investigated how they were affected by non-associative learning. We focused on the olfactory system, which contrary to visual, auditory, and somatosensory systems lacks thalamo-cortical pathways. Mitral and tufted cells, the output neurons of the olfactory bulb (OB), directly project to primary olfactory cortical areas, including the anterior olfactory nucleus (AON) and the anterior piriform cortex (aPCx) through the lateral olfactory tract (LOT) fiber bundle (Brunjes et al., 2005; Cleland and Linster, 2019). The LOT further provides input to the lateral entorhinal cortex (LEC), a multisensory structure (Tsao et al., 2013; Wu et al., 2023) situated at the intersection with the hippocampal region. The LEC, in turn, projects to the hippocampal CA1 subfield and the subiculum (SUB), a major output of the hippocampal formation (Witter et al., 2000). This olfactory cortico-hippocampal pathway is further characterized by extensive feedforward and feedback connections between the AON, aPCx, and LEC (Haberly and Price, 1978; Leitner et al., 2016; Neville and Haberly, 2003; Su et al., 2009, p. 209) on one hand and the LEC, CA1, and SUB on the other (Craig and Commins, 2007; Witter and Moser, 2006).

Non-associative learning modulates olfactory neural representations already in the OB, the first processing stage of the olfactory pathway. Brief, repeated odor experience leads to a gradual and long-lasting weakening of mitral cell odor representations in awake mice (Kato et al., 2012). In the aPCx, which has been suggested as a site of perceptual learning (Wilson and Stevenson, 2003), repeated presentation of odorants leads to short- term habituation (Wilson, 1998). Over longer time scales, neurons have been observed to change their responsiveness to odors unless they are frequently exposed to them, which has been interpreted as representational drift (Schoonover et al., 2021). How non-associative learning affects olfactory representations in the other structures of the olfactory hippocampal pathway, namely the AON, the LEC, CA1, and SUB, has not been reported.

To investigate the transformation of odorant representations across multiple brain areas along the cortico- hippocampal pathway, we used high-density neural probes (Steinmetz et al., 2021; Van Daal et al., 2021) to record more than 1400 neurons in the AON, aPCx, LEC, (ventral) CA1, and SUB of mice performing an olfactory non-associative learning paradigm, in which we presented familiar and novel odors. This paradigm allowed us to compare the encoding of chemical identity between regions and analyze at the cellular and population level how representations are shaped by experience.

## RESULTS

### Olfactory responses in the cortico-hippocampal pathway

We used a non-associative olfactory learning paradigm (Rabell et al., 2017) in which we repeatedly presented four odorants in random order to head-restrained mice (n = 16) in one daily session over four consecutive days (long-term habituation, Figure 1A and S1A). After this familiarization period, we recorded single units along the olfactory pathway (Figure 1B), specifically in the AON (n = 215), aPCx (n = 266), LEC (n = 233), CA1 (n = 474) and SUB (n = 186). We identified significant odor responses by comparing the firing rate before and after the onset of the odor with a Wilcoxon rank test (Bolding et al., 2020). In all areas investigated, we found odorant-responsive neurons (Figures 1C, 1D and S1B). The fraction of neurons that responded significantly to at least one odor was slightly larger in the AON and aPCx than in the LEC, CA1, and SUB, but overall differences were not significant (Figure 1E).

**Figure 1.**
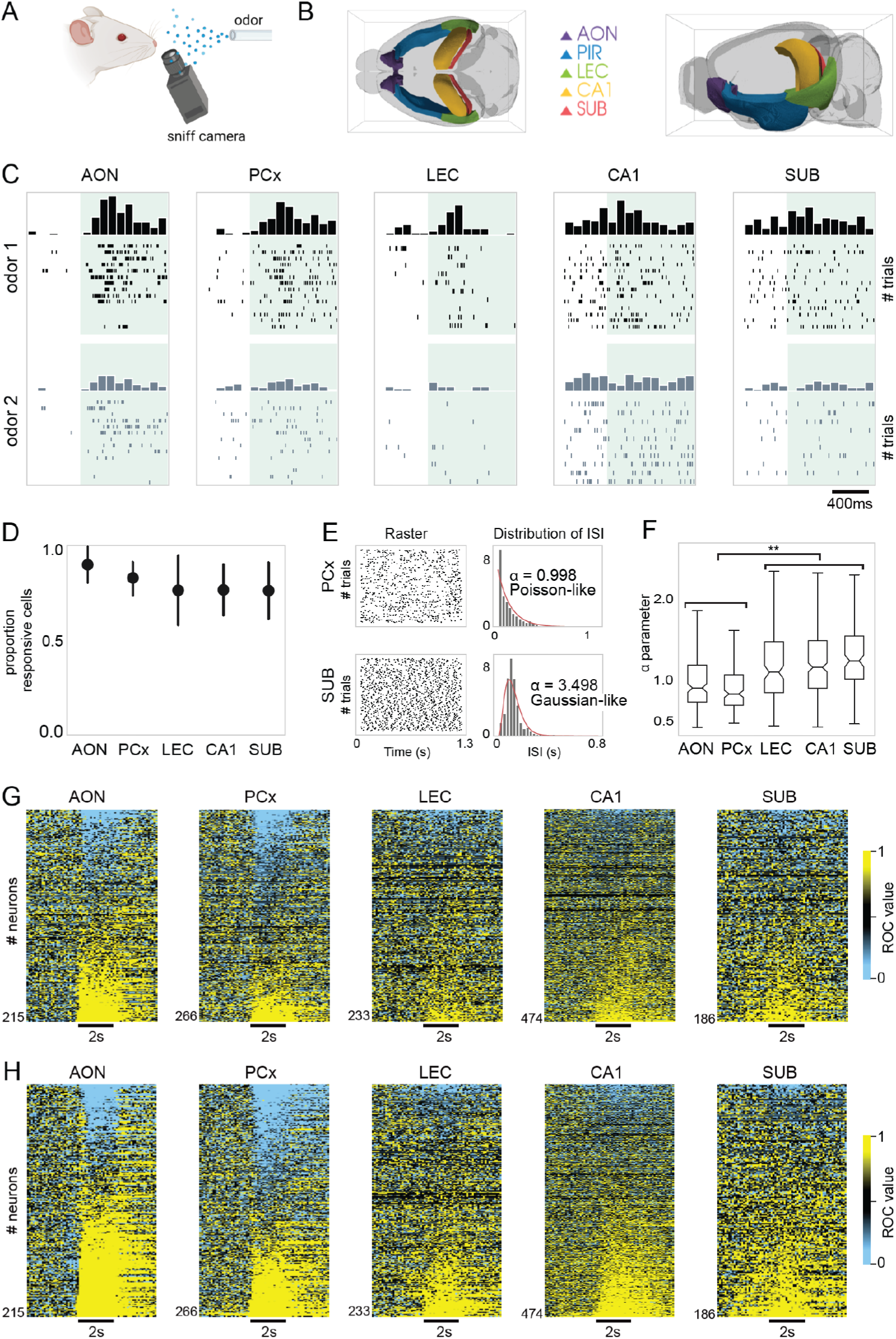
Olfactory responses in the cortico-hippocampal pathway. **(A)** Schematic representation of the experimental setup. Infrared thermography was used to measure respiration. **(B)** Illustration of recorded brain areas AON, aPCx, LEC, CA1, and SUB. **(C)** Examples of neurons responding to either of two different familiar odors in each of the five recorded areas. Green shading indicates the time window of odor delivery. **(D)** Fraction of recorded neurons per area responding to at least one odor. Odor responsiveness per area (not significant, *p>*0.3, Kruskal Wallis test). Error bars: ± standard deviation (SD) between experiments. **(E)** Raster plot during presentation of odors for two example neurons in the aPCx (top) and the SUB (bottom). Histogram of the ISI for those neurons and fitting value of gamma function (right). **(F)** Distribution of alpha parameters (*p<*0.01, Kruskal-Wallis). AON and aPCx significantly different from LEC, CA1, and SUB (^**^*p<*0.01, Mann-Whitney). (G) Responses of all recorded neurons (n=1414) to familiar odors. Yellow indicates an increase from baseline; blue indicates a decrease. Each row represents one neuron. **(H)** Responses of all recorded neurons (n=1414) to novel odors. Yellow indicates an increase from baseline; blue indicates a decrease. Each row represents one neuron.

In other sensory modalities, it has been observed that neuronal spike train statistics are different between primary sensory and multisensory neocortex. The interspike-intervals (ISI) of neurons in sensory areas follow Poisson-like statistics, whereas multisensory regions tend to exhibit more regular firing (Maimon and Assad, 2009; Shin and Moore, 2019). Fitting a gamma function to the ISI distributions of each recorded brain area reveals a similar pattern for the olfactory hippocampal pathway. Only for AON and aPCx, the shape or alpha parameters of the fitted gamma functions were close to 1, indicating a Poisson-like spiking dynamic (Figures 1F and 1G). The LEC, CA1, and SUB showed larger alpha values, consistent with more regular firing. While responses to odorants are thus found throughout the pathway, only AON and aPCx exhibit characteristics observed in sensory neocortex.

### Experience modulates odor responses across the cortico-hippocampal pathway

To investigate the impact of non-associative learning on odorant representations along the olfactory hippocampal pathway, we introduced novel stimuli randomly interleaved with familiar stimuli in our olfactory paradigm (Figure S1A). This paradigm allowed us to directly compare how chemical identity and experience (i.e., stimulus familiarity) are represented in different brain areas.

It is well established that novel odorants evoke a stereotypical increase in respiration frequency, also referred to as exploratory sniffing (Wesson et al., 2008). We used exploratory sniffing as a behavioral measure of experience. As expected, mice responded to novel stimuli with a significant increase in breathing rate (Figure S2A, *p<*0.01, paired t-test). Within one experimental session, behavioral responses to novel stimuli habituated with repeated stimulus presentation (short-term habituation, Figure S2A) such that they became indistinguishable from those to long-term habituated, familiar stimuli (*p>* 0.1, paired t-test, after three stimulus presentations).

Experience affected neuronal odorant responses in all regions studied (Figure 2A). The differential behavioral response to novel and familiar stimuli might confound the analysis of the effect of experience on sensory representations, in particular in regions in which respiration modulates spiking activity (Bathellier et al., 2008; Bolding and Franks, 2018). Therefore, we focused all further analysis on a 400ms window around the first inhalation after odor onset, when the respiration rate was not yet diverging between novel and familiar stimuli (Figures 2B and S2B; see Figure S2C-D for analysis of the entire odorant presentation period).

**Figure 2.**
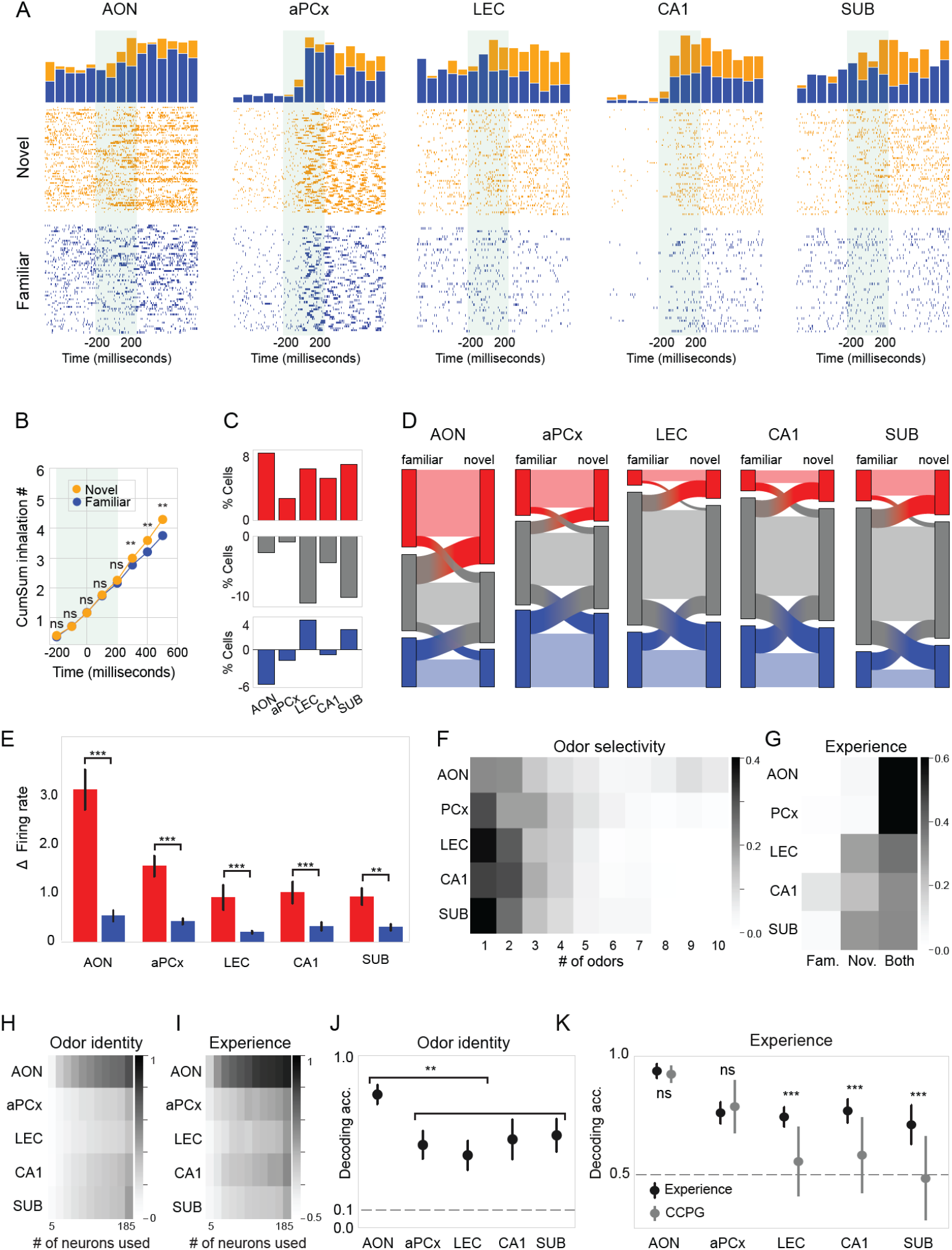
Experience modulates odor responses across the cortico-hippocampal pathway. **(A)** Example raster plots of one neuron in each brain area during the presentation of either a novel (orange) or a familiar (blue) odor. The analysis window is highlighted in green in the raster plots. The histograms represents the spike/trials per 50ms bins of time. **(B)** Cumulative inhalation counts for all experiments in the time window around odorant delivery (time point 0). All subsequent analyses were performed in a 400ms analysis window (highlighted in green), when breathing was indifferent between novel and familiar conditions (*ns*: not significant, ^*^*p<*0.05, ^**^*p<*0.01, paired t-test). **(C)** Difference in the proportion of neurons significantly excited (red) or inhibited (blue) by novel relative to familiar odors. The difference in the proportion of unresponsive neurons between novel and familiar conditions is shown in grey. **(D**) Sankey diagram showing the direction and magnitude of changes in the proportion of neurons significantly excited (red) or inhibited (blue) by familiar (left) and novel (right) odors. The direction and magnitude of changes in the proportion of unresponsive neurons is shown in grey. The width of the band represents the number of neurons in each category. Neurons with mixed response patterns were excluded from this analysis. (E) Absolute difference in firing rate between novel and familiar odors for excited (red) and inhibited (blue) neurons (^**^*p<*0.01, ^***^ *p<*0.001, Mann-Whitney U test). **(F)** Fraction of neurons with significant responses to a given number of odorants (1-10). **(G)** Fraction of neurons with significant responses to odors of either familiar, novel, or both stimulus categories. **(H)** Decoding accuracy of odor identities calculated for an increasing sample of neurons. **(I)** Decoding accuracy of experience calculated for an increasing sample of neurons. To ensure accuracy can be compared between regions, we subsampled in each region the largest common number of neurons recorded (185 neurons). **(J**) Decoding accuracy of odor identities (*p<*0.01, ANOVA). The AON significantly differed from all other regions (^**^*p<*0.01, Student’s t-test). Error bars: 95% CI of the mean. **(K)** Decoding accuracy of experience (black, *Experience*, error bars: SD) and decoding accuracy using cross-condition generalization performance (grey, *CCGP*, error bars: SD between excluded odors). The decoding accuracy was significantly different between regions (*p<*0.01, ANOVA). The AON had a significantly higher decoding accuracy than any other region (*p<*0.01, Welch t-test). The decoding accuracy was not significantly different between *CCPG* and *Experience* in AON and aPCx (ns), but significant in the other regions (***p<0.001, Wilcoxon test).

In this time window, neurons consistently responded to odorants by either increasing or decreasing their firing rate. On average less than 2.6% (±1.5%) of neurons had a mixed response pattern, such that they were excited by one odor and inhibited or by another odor. Thus, we quantified in each brain area the proportion of neurons which was i) significantly excited by odorants, ii) significantly inhibited by odorants and iii) not responsive to odorants. We found that the proportion of excited neurons was larger for novel than for familiar stimuli in all brain areas. The proportion of odor-inhibited neurons was lower for novel than for familiar odors in AON/PCx, but higher in LEC and SUB. The fraction of non-responsive cells was lower for novel than familiar stimuli in all regions (Figure 2C). The direction and magnitude of the changes in the proportion of neurons (Figure 2D) suggests that novel stimuli engaged a larger number of neurons than familiar ones. In addition to recruiting a larger number of neurons, novelty also evoked strong modulations of responses, in particular in the AON (Figure 2E) and during the first four presentations of the stimuli (S2E). The impact of novelty on response magnitude decreased towards the posterior regions.

Next, we compared the odor selectivity of neurons between regions. We determined how selectively neurons encoded different chemicals by counting the number of stimuli each neuron significantly responded to. We found marked differences between olfactory cortical areas and other regions (Figure 2F). Stimulus selectivity was lowest in the AON (mean=4.1±1.5 SD) and progressively increased from aPCx (3.0±1.0) to LEC (2.3±0.7), CA1 (2.4±0.8), and SUB (2.3±0.8). We then asked how selectively neurons encoded novelty and familiarity. We determined for each neuron whether it significantly responded to odorants of the novel, the familiar or both stimulus categories (Figure 2G). In olfactory cortical regions, both novel and familiar stimuli activated a large fraction of neurons. In the LEC, CA1, and SUB, on the other hand, a large fraction of neurons was selective just for novel stimuli.

To characterize sensory representations at the population-level, we used a linear Support Vector Machine (SVM). First, we focused on the representation of stimulus identity and decoded the chemical identity of odorants from neuronal firing rates during the analysis window across all experimental sessions (Figures 2H and 2J). The decoding accuracy was above chance level in all areas, indicating that the information about odorant identity is preserved throughout the pathway. The highest accuracy values were obtained in the AON, all other structures showed a marked decrease in decoding accuracy for odor identities in the population.

Next, we analyzed the encoding of experience in the neuronal population from each region by training a SVM to classify novel vs. familiar trials. We found the highest decoding accuracy of experience in the AON, while all other regions showed similar, lower levels of accuracy (Figure 2I and 2K). Overall, the neural representation in the AON thus showed the highest decoding accuracy for both odor identity and experience.

Since we had found earlier that odor identities could be decoded from the neural data, the SVM might recover the information about experience solely based on odor identity. To address this potential confound, we compared the decoding accuracy of novel versus familiar odors with the decoding accuracy using Cross-Control Generalization Performance (CCGP, Bernardi et al., 2020). In this framework, the classifier is trained on a subset of odors while leaving one odor out for testing. This approach allows us to assess how well the classifier can generalize the information of experience across different odors. High decoding accuracy with CCPG suggests the classification performance is independent of odor identity. If decoding accuracy with CCPG is low, the SVM was able to classify trials into novel and familiar based on odor identities. We found that contrary to AON and aPCx, decoding performance in the LEC-Hpc did not generalize well across odors, and in SUB the information was not generalized at all. This suggests that while early olfactory regions encode novelty in a relatively odor-independent manner, the LEC-HPc regions represent experience in a more stimulus-specific way.

### A small subpopulation of neurons in the aPCx is tuned to familiar stimuli

Non-associative learning creates implicit memories of stimuli in the environment. To investigate how this memory is reflected in the sensory representations, we compared the encoding of odor identities between novel and familiar conditions. Using an SVM to decode odorant identity separately for novel and familiar odorants, we found that experience impacts the representation of stimulus identity (Figure 3). Specifically, the aPCx showed much higher decoding accuracy for familiar odors than novel ones (Figure 3A). This might suggest the population of aPCx neurons is particularly tuned to familiar odorants. To investigate this further, we calculated the mutual information between neuronal firing and odorant identity, separately for novel and familiar conditions (Figure 3B). This measure indicates how much knowing the firing rates reduces the uncertainty about the identity of odorants. Surprisingly, mutual information was similar for novel and familiar conditions in the aPCx indicating no specific tuning of the whole population to familiar odorants. But how could the SVM then have achieved a much higher accuracy in decoding the identity of familiar odorants compared to novel ones (Figure 3A) ?

**Figure 3.**
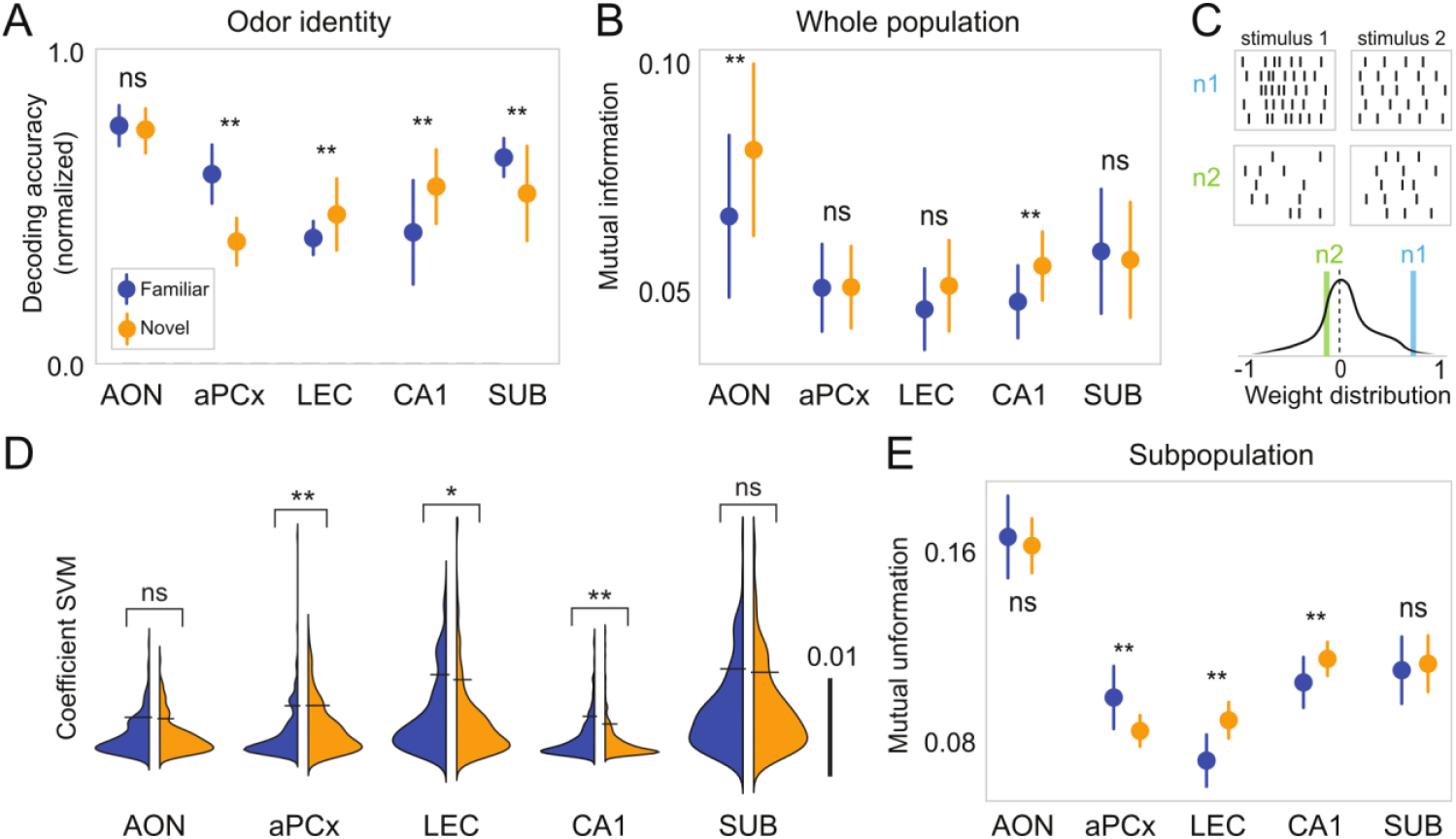
A subpopulation of neurons in the aPCx is tuned to familiar stimuli. **(A**) Accuracy of decoding the chemical identity of familiar and novel odors (^*^*p<*0.05, ^**^*p<*0.01, paired t-test). Error bars: 95% CI of the mean. **(B)** Mutual information (MI) between odor identities and neuronal responses (^**^*p<*0.01, paired t-test). Error bars: 95% CI of the mean. **(C)** Top: Simplified classification scheme in which two neurons contribute differently to decoding stimulus identity. The weight vector is orthogonal to the hyperplane separating the data. Bottom: Distribution of weights of all neurons. Neuron 1 (n1, blue) has a large weight, neuron 2 (n2, green) has a low weight in encoding stimulus identity. **(D)** Distribution of weights (absolute values) of the SVMs decoding the identity of either novel or familiar odorants. Significant differences were observed in aPCx, CA1, and LEC (*ns*: not significant, ^*^*p<*0.05, ^**^*p<*0.01, Wilcoxon rank- sum test). **(E)** MI of outlier subpopulations (*ns*: not significant, ^**^*p<*0.01, Welch’s t-test). Error bars: 95% CI of the mean.

During the training of the linear SVM, each neuron receives a weight that determines its relative contribution to the classification decision (Figure 3C). Importantly, a relatively small number of neurons with a large weight can have a decisive impact on classifier performance, yet they might not affect measures such as mutual information. When examining the weight distributions of SVMs for encoding familiar and novel odor identity, we found that the weights for decoding familiar odorants in the aPCx had a long tail of outliers (Figure 3D, example neurons shown in Figures S3 and S4). We calculated the mutual information for outlier subpopulations across all brain areas, corresponding to an average of 5.2% (± 0.8 SD) of the data for each brain area (Figure 3E). The mutual information of the outlier population now captured the differences in the accuracy of encoding familiar and novel odorants observed in the SVM (Figure 3A), in particular in the aPCx but also the LEC and CA1 regions. And while the effect size revealed through the subpopulation analysis in Figure 3E does not fully account for the difference observed in Figure 3A, it nonetheless highlights a consistent trend between multiple regions.

Collectively, these results suggest the presence of a subpopulation of neurons in the aPCx that are particularly tuned to odorants when they are familiar, as well as a subpopulation of neurons in the LEC and CA1 tuned to novel odorants. This analysis highlights that long-term, non-associative learning can refine the tuning of small fractions of neurons.

### The representations of odor identity and experience become independent from each other along the cortico-hippocampal pathway

The presence of neuronal subpopulations specifically tuned to familiar odors in aPCx and to novel odors in LEC and CA1 (Figures 3A and 3E) raises the general question of how much each neuron contributes to encoding individual stimulus features. Do separate groups of neurons represent different stimulus features, or do all neurons form a continuous distribution for “multiplexed” encoding of odorant identity and experience?

To address this question, we analyzed the weights of the SVM trained to classify odorant identity. We did this separately for novel and familiar odors. If the same neurons which were good at classifying the chemical identity of the familiar odors were also good at identifying the chemical identity of the novel odors, their weights should be correlated. We found that SVM weights were increasingly correlated across the olfactory pathway from AON to SUB (Figure 4A). This suggests that neurons specialized in encoding chemical identity, independent of experience.

**Figure 4.**
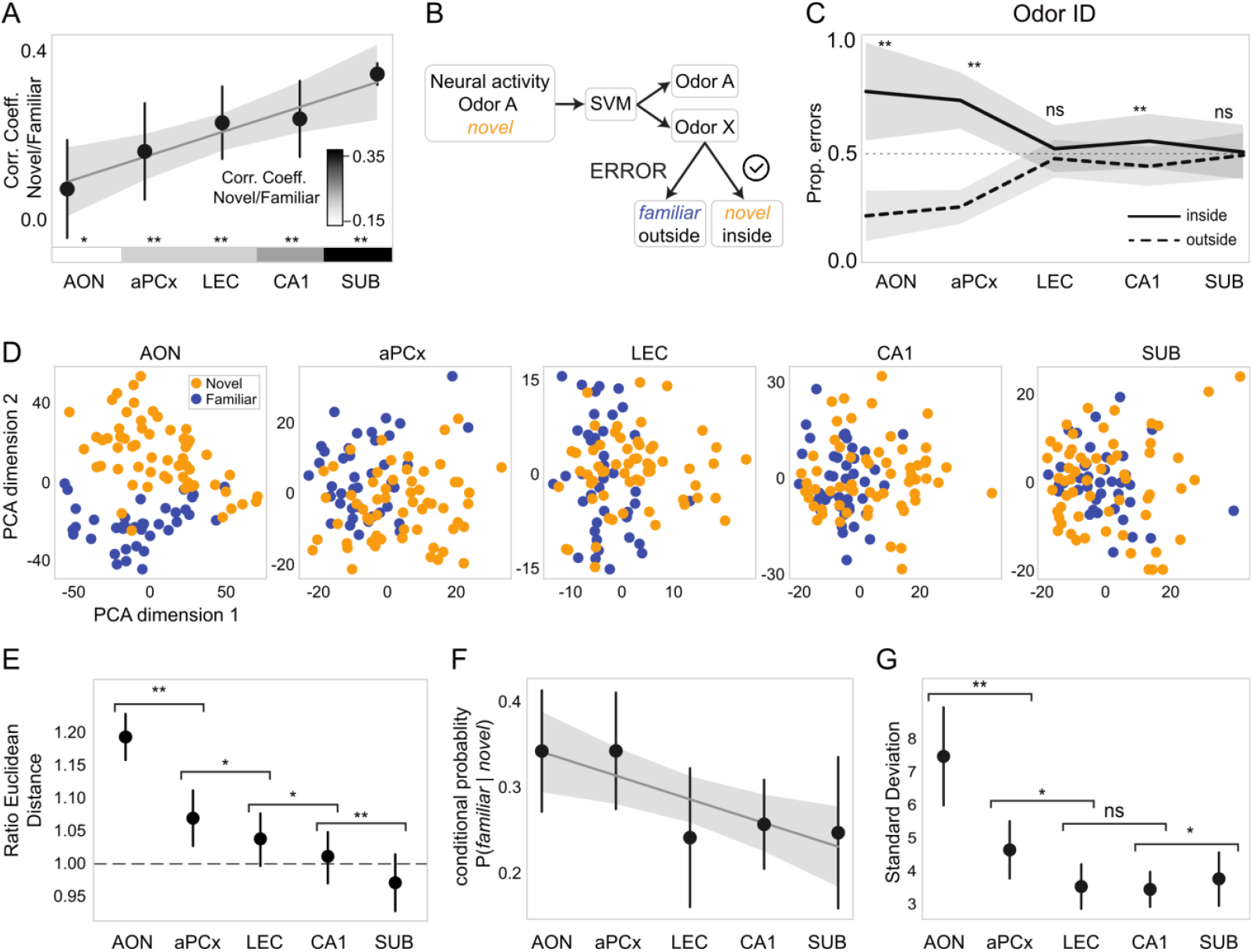
The representation of identity and experience become independent from each other along the cortico- hippocampal pathway. **(A)** Pearson correlation coefficients between SVM weights for decoding familiar and novel odor identity (^*^*p<*0.01, ^**^*p<*0.01). Error bars: Standard error of the mean (SEM). **(B)** Schematic representation of the methodology to calculate error rate inside and outside the experience category (novel vs. familiar) with the example of a novel odor A. **(C)** Proportion of errors per region. Solid and dashed lines represent the proportion of errors inside and outside the same experience category, respectively. (*ns*: not significant, ^**^*p<*0.01, Wilcoxon rank-sum test). Error bars: SD. **(D)** Position of odors in the neural space for the 10 trials of four familiar odors (blue) and six novels (orange). The first two components of the PCA were used. Percentage explained by the first two dimensions: AON 31%, aPCx 15%, LEC 12%, CA1 15%, and SUB 28%. **(E)** Normalized Euclidean distance of principal components between novel and familiar odors divided by the average distance between all trials, pooled irrespective of experience (*p<*0.01, Kruskal- Wallis). Post-hoc pairwise comparison revealed significant differences between all regions (^*^*p<*0.05, ^**^*p<*0.01, Mann- Whitney). Error bars: 95% CI of the mean. **(F)** Probability of responding to one or more familiar odors if a neuron responded to at least one novel odor (*p<*0.001, Kruskal-Wallis). AON and aPCx significantly different from LEC, CA1, and SUB (*p<*0.01, Mann-Whitney). Error bars: 95% CI of the mean. **(G)** Standard deviation of the firing rate in response to olfactory stimuli (*p<*0.001, Kruskal Wallis test). Response variability progressively decreases from AON to SUB (*ns*: not significant, ^*^*p<*0.01, ^**^*p<*0.01, Mann-Whitney). Error bars: 95% CI of the mean.

If this was indeed the case, it should also be reflected in the errors made by the SVM. If a neuronal population is specialized in encoding chemical identity irrespective of experience, a classification error will correspond to mistaking an odor for any other odor. On the other hand, if separate populations encoded familiar and novel odors, a SVM should mostly mistake familiar odors for other familiar odors and novel odors for other novel odors (Figure 4B). Analyzing the SVM errors for each brain area, we found a shift from making errors within the same category (i.e., the decoder mistakes a familiar odor for another familiar odor or a novel odor for another novel odor) to mistakes being made interchangeably between novel and familiar categories (Figure 4C).

The finding that encoding stimulus identity becomes independent of experience in multisensory areas is further supported by analysis of neuronal firing at the population level. We used principal component analysis (PCA) to investigate the position of novel and familiar trials in neural space. Notably, the representations of stimulus identity were separated between novel and familiar stimuli in AON and aPCx, while they became increasingly similar in multisensory areas (Figure 4D).

To quantify this effect, we calculated the Euclidean distance between novel and familiar trials on the one hand and between all trials irrespective of experience on the other. In AON, aPCx and LEC, the distance between novel and familiar trials was significantly higher than when not taking experience into account. Conversely, in the SUB, novel and familiar trials were significantly closer than the average distance between all trials (Figure S6A). The ratio of the two Euclidean distance measures revealed a significant, progressive decrease in distance between familiar and novel trials along the pathway (Figure 4E).

Given the representation of odorant identity became progressively independent from the representation of experience, we then asked if there was a distinct population of neurons encoding experience, independent of odorant identity. The SVM could decode experience in the multisensory areas with high accuracy (Figure 2I), but we found little evidence for the emergence of neurons that specifically responded in a stimulus-invariant manner to all novel or all familiar stimuli (novel 1/1414s and familiar 2/1414; Grubbs test confirmed stimulus-invariant neurons were not rare outliers of the distribution; see methods). Instead, we observed an increasing odor selectivity in multisensory areas (Figure 2D) and the emergence of a population of neurons responding significantly to specific odors only when they were novel (Figure 2E). Across the olfactory-hippocampal pathway, if a neuron responded to at least one novel odor, it had a decreasing probability of also responding to a familiar odor (Figure 4F). These observations imply that the number of odor-responsive neurons reflects experience. In any given trial, the more neurons were activated by a stimulus, the more likely the stimulus was novel.

In conclusion, we describe the transformation of sensory representations from olfactory cortical regions to lateral entorhinal-hippocampal areas, during which the representations of odorant identity and experience became increasingly independent from each other. This transformation of sensory representations coincided with increasing response reliability, as we observed that the variance of neuronal responses progressively decreased from AON to LEC-Hpc (Figure 4G).

## DISCUSSION

In this study we carried out extracellular recordings in awake, head-restrained mice performing a non- associative olfactory learning paradigm to characterize the transformation of olfactory representations from olfactory cortical regions to entorhinal hippocampal areas. We analyzed neural responses to different novel and familiar olfactory stimuli in the AON, aPCx, LEC, CA1, and SUB to determine how odorant identity and odorant experience were reflected in each brain area. We found that both odor identity and olfactory experience were represented in every region, but representations were transformed between brain areas along the pathway. In the AON, neurons were broadly tuned to different chemicals, and their responses were modulated by experience such that novel stimuli caused larger increases in firing rates than familiar stimuli. From the AON to hippocampal structures, the selectivity of neurons for specific odorants increased, concurrent with the development of population-level odor representations, which became independent of novelty and familiarity. Consistent with this, we observed the emergence of specific populations of neurons tuned to select familiar odorants in aPCx and novel odorants in LEC-Hpc structures. These results suggest a progressive separation of the representations for identity and experience from AON to SUB.

The very first cortical processing stage, the AON, consistently outperformed all other brain areas, including the aPCx, in decoding accuracy for both stimulus features. Although the AON is the earliest cortical brain region involved in olfaction, it has thus far received remarkably little attention. This is in part because past research has focused on the aPCx, which is widely believed to carry the information for determining odorant identity (Bolding and Franks, 2017; Kadohisa and Wilson, 2006; Miura et al., 2012; Stettler and Axel, 2009). Further research is needed to clarify the relative contribution of AON and aPCx in odorant identification. Moreover, we acknowledge the lack of recordings from posterior PCx (pPCx) in our study. The pPCx is anatomically positioned between AON, aPCx and the LEC-Hpc areas and is often considered as associational structure(Wang et al., 2020). Investigating how experience affects odor responses in the pPCx will provide additional insights into the transformation of neural representations in the olfactory cortico-hippocampal network.

The finding that the LEC, CA1, and SUB represent the identity of odorants after they have been familiarized by passive exposure over several days might be considered surprising since nondeclarative learning is commonly viewed to be independent of medial temporal lobe structures (Squire et al., 2021, 2004). However, a specific projection from the CA1 to the AON has recently been shown to be critical for spatiotemporal odor memory in a nondeclarative behavioral paradigm (Aqrabawi and Kim, 2020). Accordingly, the representation of specific odorant chemicals we observed might contribute to the formation of contextual odor memories in CA1 and structures of the LEC-Hpc areas.

The analysis of the representation of experience in the LEC, CA1, and SUB offers additional important insights. Novel stimuli activated more neurons than familiar stimuli in the LEC, CA1, and SUB, which is in line with earlier descriptions of novelty-related activity in the hippocampus (Rolls et al., 1993). However, we did not find evidence for an abstract representation of novelty by stimulus-invariant “*novelty*” neurons. Consistent with the fact that hippocampal lesions do not disrupt novelty detection (Aqrabawi and Kim, 2020; Honey et al., 1998), our results thus offer a new perspective on the widely accepted interpretation that the hippocampus conveys a novelty signal to other brain structures, which broadly enhances attention and memory (Lisman and Grace, 2005; Ranganath and Rainer, 2003). Of note, previous studies had limited temporal or cellular resolution (Bunzeck and Düzel, 2006; Zhu et al., 1995), often used few novel stimuli, and did not report on stimulus-invariance (Xiang and Brown, 1998). Rather than encoding abstract stimulus novelty, our data suggests that the highly stimulus-selective neurons in LEC, CA1, and SUB represent stimulus identity, but an attentional gating mechanism causes their preferential activation when stimuli are novel. Further research is required to validate this hypothesis and to clarify the stimulus-invariance of individual neurons in response to novel stimuli from other sensory modalities and complex, multimodal stimuli, such as novel environments.

How experience affects sensory cortical representations has thus far mainly been studied in the visual system. Compared to familiar stimuli, novel stimuli induce larger activation in the visual cortex (Garrett et al., 2020; Homann et al., 2022), which can be explained by plasticity mechanisms inherent to neocortical circuits (Aitken et al., 2024). We found the same effect of stimulus novelty on responses in the AON, which is remarkable, given the anatomical differences between the AON, a 2-layered archeocortical structure, and the 6-layered neocortex. It will be interesting to further investigate how novelty is processed in circuits with diverging architectures.

In conclusion, we propose a model (Figure 5) in which olfactory cortical areas represent stimulus identity with broadly tuned neurons, whose response magnitude is modulated by experience. This representation is transformed such that stimulus identity becomes encoded by more narrowly tuned neurons and experience by the number of active neurons. Future research involving simultaneous recordings along the cortico-hippocampal pathway will provide further insights into the processing of novel and familiar olfactory stimuli. While technically more challenging, simultaneous recordings can help to establish the temporal order of activation of different brain areas in order to resolve how the intricate network of feedforward and feedback connections supports encoding of stimulus identity and experience. Moreover, further work is needed to address the question if our findings extend to declarative paradigms, such as operant novelty detection tasks.

**Figure 5.**
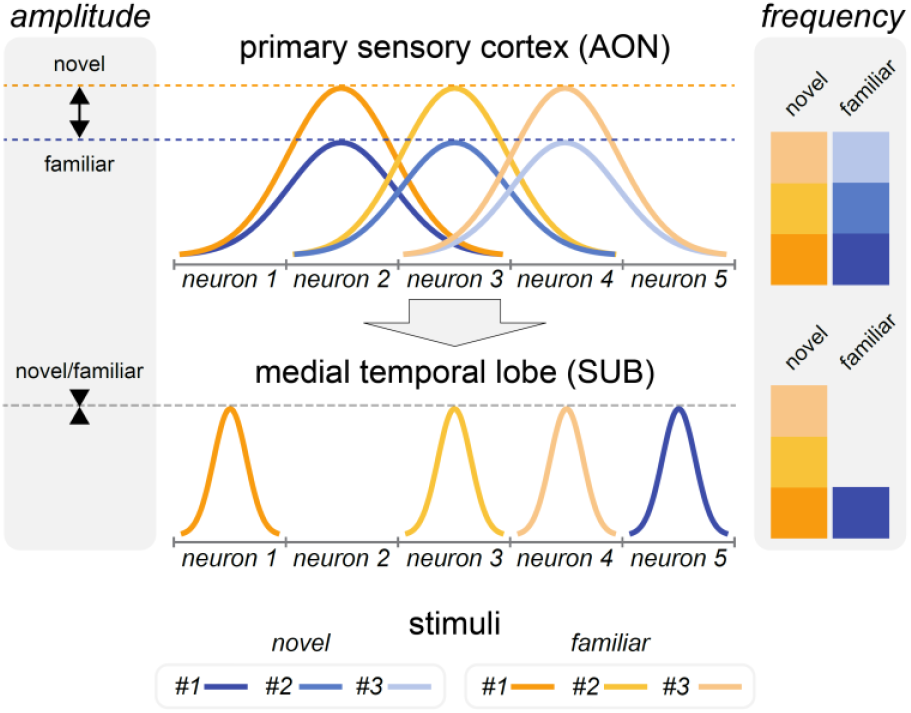
Proposed mechanism for the transformation of olfactory representations. Illustration showing the responses of 5 neurons in AON (top) and SUB (bottom) to either of 3 novel (orange) or familiar (blue) odorants. Novel stimuli evoked larger responses than familiar stimuli in the AON, whereas in hippocampal areas, novelty was reflected by the number of responsive neurons. Moreover, the stimulus selectivity decreases from AON to SUB (narrowing of response curves).

## ACKNOWLEDGMENTS

This work was supported by Interne Fondsen KU Leuven C14/21/111 (SH), Fonds Wetenschappelijk Onderzoek-Vlaanderen (FWO-Flanders) fellowships 1135122N (ES) and 1272222N (CA), and grant G097022N (SH).

## MATERIALS AND METHODS

### Experimental model

Experiments were approved by KU Leuven Animal Ethics Committee under protocol P067/2017 and performed in accordance with the Federation of European Laboratory Animal Science Associations.

We used 13 male and 3 female mice between 18 and 30 weeks old, C57BL/6J (from KU Leuven). Animals were housed on a 12 h dark/12 h light cycle (dark from 07:00 to 19:00). Before the experiment, they were housed in groups of 2 to 4 mice until adult age, and at the start of the experiment, they were individually housed in GM500 cages (391 × 199 × 160 mm) with food and water *ad libitum*. Before starting behavioral tasks, animals were habituated to head-restrain for two days. All animals performed their task around the same time each day.

### Surgical procedures

All mice were surgically implanted with a stainless-steel head plate to allow for head-restraining. Animals were anesthetized with ketamine and medetomidine mixture (60mg/Kg and 0.5mg/Kg of body weight intraperitoneally), and the depth of anesthesia was checked by monitoring tail pinch response, whisking and eye reflexes (as regularly throughout surgeries). If necessary, a booster shot of half the initial amount of anesthesia was given during the surgery. The head was shaved and then washed down with a series of Betadine (Meda, cat. no. 2990-257). The Xylocaine 5% (wt/wt) (Aspen, cat. no. 0137-398) was administered subcutaneously at a dose of 13 mg/Kg. Duratears (Alcon, cat. no. 0037-820) was applied to avoid dehydration of the eyes. Then mice were placed on a heating pad at 37°C (Harvard apparatus) in a stereotaxic frame (Narishige Instruments) using ear bars. An incision was made longitudinally along the top of the head, and a window of 2cm^2^ was cut out to expose the underlying skull. Less than 50μL of Vet Bond (3M) was applied at the border between the skin and the skull to prevent irritation and scratching. A scalpel was used to remove connective tissue. The skull was dried using an air-blow device for 1 minute. The site of the craniotomy was then marked using a scalpel. Using a dental drill, ridges were created in the bone to increase the surface area of the skull for more adhesion. Dental cement (acrylic) was applied to glue the custom-made titanium head plate to the bone while leaving the site of the craniotomy free. The cement was let to dry for 10 minutes. The craniotomy was made in the shape of a circular hole of 1.5-2mm. A durotomy was made using a small precision scissor and a tweezer.

Subsequent steps following the craniotomy were performed according to whether recordings were conducted under acute or chronic conditions. For acute recordings, we used Neuropixels 1.0 probes (n=6); for chronic recordings, we used Neuropixels 2.0 probes (n=10). Before brain insertion, all probes were marked with a fluorescent dye (diI, Life Technologies). In animals designated for acute recordings, the craniotomy was covered with artificial dura to keep the exposed cortex healthy. Then, the craniotomy was closed with silicone (Kwik-Sil, World Precision Instruments). In animals designated for chronic recordings, probes were inserted using a micromanipulator at 20 μm/s to 60μm/s. Bone wax (Surgical Specialties, cat. No. 901) was applied over the probe insertion site to maintain a closed environment around the craniotomy site. Light curable dental cement (SDI Wave, cat. no. 7510203) was applied to secure the probe to the skull by curation with blue light.

At the end of each surgery, atipamezole (antisedan, 0.5mg/kg) was administered IP, immediately followed by the analgesic Metacam at 5-10 mg/kg. Animals were continually monitored until they regained consciousness.

### Probes and coordinates

| Type | Region | n= | Probe | AP (mm) | LM (mm) | Depth | Angle (°) |
| --- | --- | --- | --- | --- | --- | --- | --- |
| Acute | AON | 3 | 1-shank | 2.55-2.96 | 0.81-1.50 | 2.50-3.10 | 0-10 (latero-medial) |
|  | aPCx | 3 | 1-shank | 1.00-2.34 | 1.25-1.95 | 2.75-4.00 |  |
| Chronic | AON | 2* | 4-shank | Shank 0: 2.90 | 0.80 | 3.00 |  |
|  |  |  |  | Shank 1: 2.70 |  |  |  |
|  |  |  |  | Shank 2: 2.50 |  |  |  |
|  |  |  |  | Shank 3: 2.35 |  |  |  |
|  | aPCx | 4* | 4-shank | Shank 0: 2.10-2.40 | 1.35-1.95 | 3.65-4.90 | 10 (anterior-posterior), 0-10 (latero-medial) |
|  |  |  |  | Shank 1: 1.80-2.25 |  |  |  |
|  |  |  |  | Shank 2: 1.70-2.10 |  |  |  |
|  |  |  |  | Shank 3: 1.50-1.94 |  |  |  |
|  | LEC & Hippocampus | 5 | 4-shank | Shank 0: -3.535 | 2.75 | 4.50-5.00 | 15 (latero-medial) |
|  |  |  |  | Shank 1: -3.285 |  |  |  |
|  |  |  |  | Shank 2: -3.035 |  |  |  |
|  |  |  |  | Shank 3: -2.785 |  |  |  |

\*one animal yielded units in both AON and aPCx.

### Electrophysiological recordings

During acute recordings, the silicone seal covering the craniotomy was removed, and the probe was connected to the recording hardware and inserted using a micromanipulator. For chronic recordings, implanted probes were connected to the recording hardware. Setups were denoised with grounded metal structures attached to the recording system. Raw data was acquired using SpikeGLX software on a computer specifically dedicated to this use. Experimental events were acquired at 3 kHz by analog inputs on SpikeGLX. Respiration was monitored with a thermal camera (FLIR A325sc with 50μm lens, 60fps, 320×240 pixels) directed to the nostrils of the animal to capture respiration.

### Odorants

The stimulus set consisted of 28 single-molecule substances (Sigma-Aldrich) commonly used in olfaction research. Odors were benzyl acetate, nonadienal, isoamyl acetate, heptanal, ethyldecanoate, linalyl formate, trimethylpyrazine, geraniol, geranyl acetate, salicylaldehyde, alpha-pheleandrene, thiophane, phenylethyl alcohol, dimethylpyrazine, limonene, carvone, octanoic acid, acetic acid, menthol, eugenol, benzaldehyde, ethyl methyl phenylglycidate, anisole, cinnamaldehyde, ethyl valerate and pentenoic acid. Each odor was dissolved in mineral oil to reach a final vapor pressure of 1Pa. Odors were delivered with a custom-made, constant air flow olfactometer, as described previously (Rabell et al., 2017).

### Non-associative olfactory learning paradigm

All experiments were performed inside a sound and light isolated box (75*75*75 cm3) with constant exhaust to ensure rapid clearance of odorants. After recovery from the surgery (minimum 3 days), mice underwent habituation to head-restraining in the setup for one to three days. The paradigm was then performed essentially as described previously (Morrens et al., 2020). First, mice were familiarized with 4 odorants for 4 consecutive days. On subsequent days, novel odorants were introduced randomly interleaved with the previously familiarized odorants. All odors were delivered for 2 seconds and presented 10 times within one session. In recordings performed in AON and aPCx, a total of 8 novel odors were presented per session; in all other recording sessions, we used 6 novel odorants. To ensure that results were comparable between regions, we restricted our analysis to the first 6 novel stimuli presented. AON/aPCx recording sessions started with 4 presentations of familiar odorants, after which novel and familiar stimuli were presented in random order. In all other sessions, the order of odorant delivery was fully randomized. Two recording sessions were excluded from the analysis because 3 odorants which had been introduced as novel on the previous day, were mistakenly presented again.

### Histology

Upon completion of experiments, mice received an overdose of ketamine/medetomidine, exsanguinated with PBS, and perfused with 4% paraformaldehyde. Brains were cut into 80μm coronal sections on a vibratome. Brain slices were mounted on glass slides using a VectaShield medium and analyzed using a confocal microscope to reconstruct anatomical recording locations based on the DiI label (Life Technologies), the probe’s apparent trajectory, and the implantation’s depth.

## QUANTIFICATION AND STATISTICAL ANALYSIS

### Processing of respiration signals

Videos recorded with the thermal camera were processed as described previously to extract inhalation onsets (Mutlu et al., 2018).

### Preprocessing of the neural data

Electrophysiology data was preprocessed as described previously (Van Daal et al., 2021), using a combination of CatGT and kilosort 2.5. Following the automatic clustering, neurons were manually curated using phy2 to avoid multi-unit clusters, drifting neurons, or clusters with no refractory period.

### auROC odor responses

We used the area under the receiver operating characteristic curve (auROC) to compute the discriminability of the response between baseline and odor presentation. Spiking activity was first divided into 100ms bins. The baseline firing before odor delivery (the first 2s of each trial) was then compared with each bin afterward.

### Responsiveness to odor and modulation by experience

Significant odorant responses were identified by comparing baseline firing rates before odorant delivery during the first 10 presentations of a given odor to the firing rate during the analysis window (Figure 2B) with a Wilcoxon rank test. A neuron was considered responsive to an odor if the test result was statistically significant. A similar approach was used to determine significant modulation by experience, performing the Wilcoxon rank test separately for novel and familiar stimuli. Neurons significant for both novel and familiar conditions were indicated as responding to both.

### Gamma fit

We used a methodology described previously (Maimon and Assad, 2009). In brief, the Inter-Spike Interval (ISI) of selected neurons was computed during the presentation of odors. The histogram distribution of the ISI was then fitted using a gamma function. Alpha parameters close to 1 signal a Poisson-like spiking dynamic.

### Population decoding

Decoding was done using a linear Support Vector Machine (Kramer, 2016) to determine the identity of odorants (multi-class) or level of experience (binary class) using the firing rate of neurons in the analysis window (Figure 2B). Neurons from all sessions were concatenated. To allow for comparing the accuracy between regions, we subsampled the same number of neurons for each region (185 neurons). During 100 iterations, neurons were randomly chosen from the entire pool of neurons. Subsequently, neurons were divided into distinct sets for training and testing the SVM for each iteration. In order to mitigate the issue of overfitting, the Leave-1-out approach was used. Since sample sizes differed between regions, each iteration of the leave- one-out approach was averaged across different pools of decoded subpopulations. Thus, the variation (95%CI or SD) represents the fluctuation of performance of the SVM in trial-to-trial differences.

As there were more novel than familiar stimuli in our experimental paradigm, we took measures to balance the dataset for the SVM. For decoding experience, we used the first 40 trials of familiar and novel stimuli, respectively. For decoding odorant identity of novel and familiar stimuli separately, we used a Min-Max normalization (1) to balance different chance levels of decoding novel and familiar stimuli (Figure 3). (*DA*_*normed*_: normalized decoding accuracy, *Pc*: chance level chance of the stimulus).

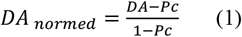

### Amplitude of the response

The difference in response amplitude to novel and familiar odors was computed for each neuron and was defined as the absolute value of the mean response to novel odor minus the mean response to familiar odor.

### Sankey diagram

For each neuron, baseline firing rate was calculated during the pre-odor period and compared to the firing rate during the analysis window using a Wilcoxon rank-sum test. Significance was assessed separately for novel and familiar odors. Neurons showing a significant change were classified as *excited* if firing increased relative to baseline and *inhibited* if firing decreased.

### Mutual Information

Mutual information was calculated using the scikit learn library (Kramer, 2016) using a log base 2 (resulting in Shannon units – or bits). This allows for measuring the dependencies between two variables based on the methodologies described previously (Kraskov et al., 2004), which gives an entropy estimation from k-nearest neighbors distances (k=3). There is one discrete variable (odor identities) and one continuous (firing rate over consecutive trials). The function *mutual_info_classif* was used to quantify the amount of information recoverable on odor identities from the firing rate of cells. The mutual information was computed ten times for each cell-odor pair of one odor versus another odor (novel or familiar). This approach was repeated for the subpopulation analysis (Figure 3E).

### Probability of responding to familiar odors

*Fam*_*Normed*_ and *Nov*_*Normed*_ refer to the normalized number of familiar and novel odors, respectively, to which a neuron significantly responded.

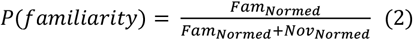

For each neuron responding to at least one novel odor, (2) was applied to obtain *P*(*familiarity∣novelty*)

### Grubbs test and novelty decoding

Using formula (2), *P*(*familiarity*) and *P*(*novelty*) were computed for each neuron responding to at least one novel or one familiar odor. A value of 1 for *P*(*novelty*) indicates that the neuron is responding only to novel odor. A Grubb test was then used to evaluate whether those values indicated a particular outlier outside the normal range of the distribution.

### Weight of the SVM

Weights of the SVM were extracted for the identity decoding of familiar and novel odors. The absolute value of those weights was used for the following analysis. The Pearson correlation for those weights was computed in Figure 4A. For the subpopulation analysis, neurons that had weights two times the standard deviation above the mean were counted (for familiar odors, AON=12, aPCx =11, LEC=13, SUB=8, CA1=24 and for novel odors, AON=15, aPCx =15, LEC=14, SUB=8, CA1=22) and used to compute the MI (Figure 3E).

### Confusion matrix and proportion of errors inside/outside level of experience

The confusion matrix for the decoding of all odors was computed. The mean error inside the same category (novel odors are mistaken for other novel odors; familiar for other familiar odors) and outside the category (novel odors are mistaken for familiar odors; familiar for novel odors) were then assessed. The proportion of errors made inside or outside the category was then quantified. The variance reflects the variability between different test/training sets.

### Distance in the neural space

The spike rate of each neuron in the analysis window was extracted for the first ten presentations of familiar and novel odors. They were reduced into a 2D space using a PCA. The PCA space was normalized using a Min-Max method (3) per region and for each dimension. The distance between trials was evaluated using Euclidean distance in 2D. The inter-experience distance was measured between the familiar odor trials and the first six novel odors trials. The ratio of Figure 4G was computed by dividing the inter-experience distance by the inter-trials distance.

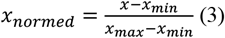

