## Supplementary file for "Experience shapes the transformation of olfactory representations along the cortico-hippocampal pathway"

### SUPPLEMENTAL INFORMATION

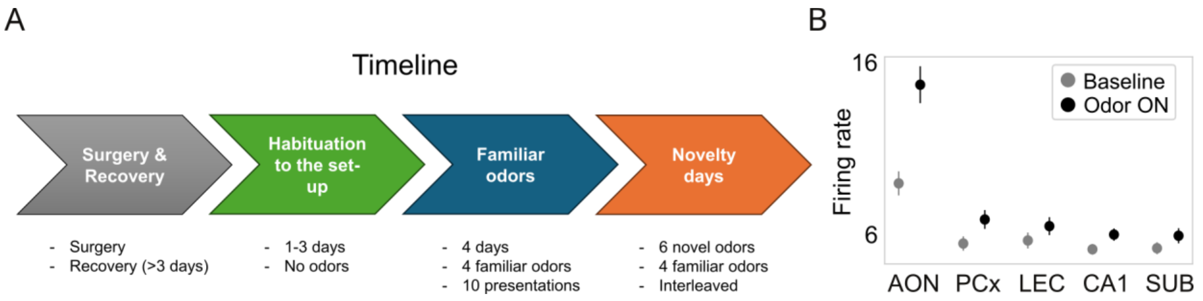


**Figure S1. Experimental protocol and response to odors. (A)** Timeline of the experimental protocol. **(B)** Average firing rate of neurons at baseline and during the first presentation of an odor (2s). Values are significantly different in each region (Wilcoxon paired test, $p<0.05$). Error bars: Standard error of the mean (SEM).

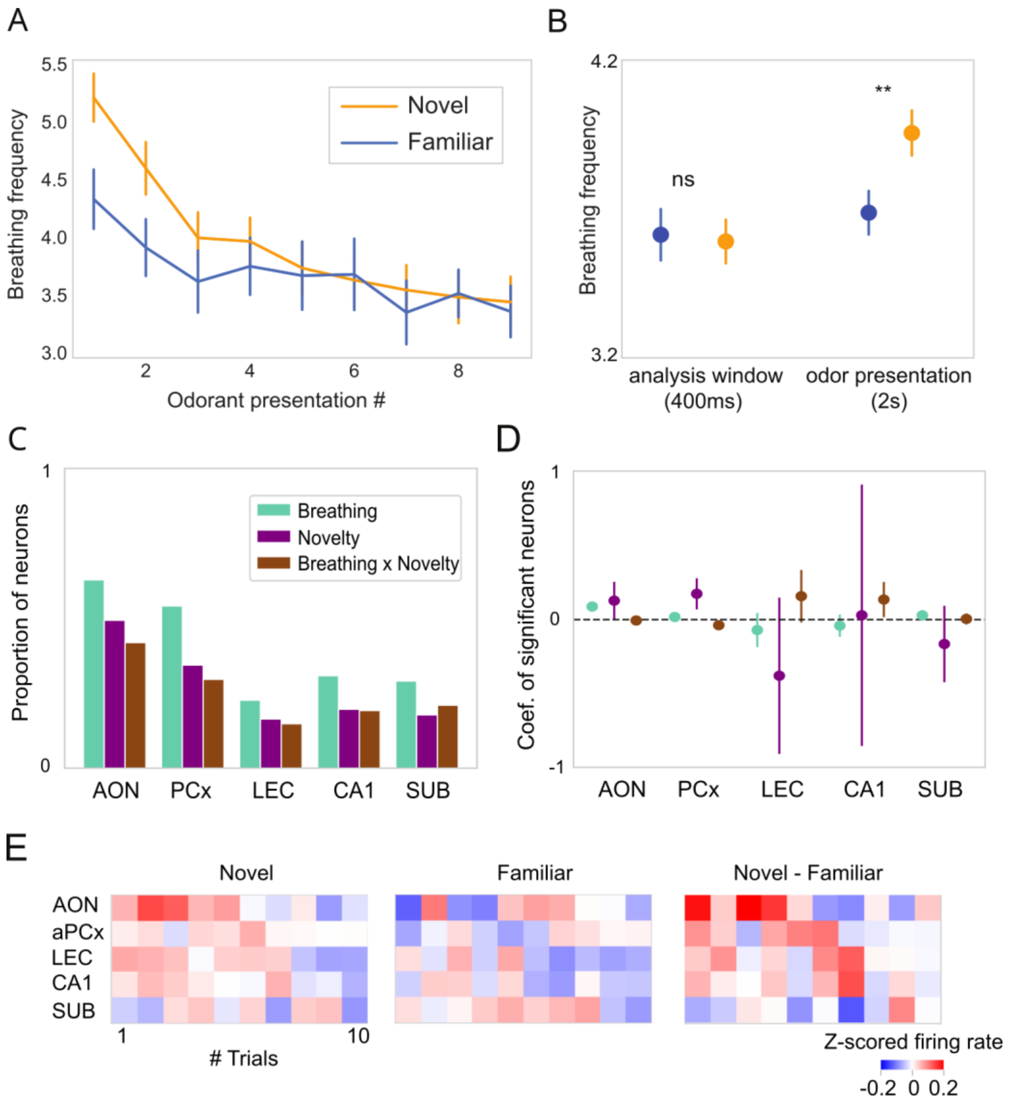


**Figure S2. Exploratory sniffing behavior and response to experience. (A)** Breathing rate in response to novel (orange) or familiar (blue) odorants. Repeated presentation reduces responses (short-term habituation). Error bars: Standard error of the mean (SEM). **(B)** The average breathing rate is not significantly different between the novel and familiar odorants during the analysis window (400ms, Figure 2B), but when taking the entire period of olfactory stimulation (2 seconds) into account (*ns*: not significant, $^{**}p<0.01$, Wilcoxon rank-sum test). Error bars: Standard error of the mean (SEM). (C) General Linear Model for entire odor presentation window (2s). Proportion of neurons significantly affected by each factor ($p<0.05$, two-tailed t-test) (D) Distribution of significant coefficients for novelty, breathing and the interception term. Error bars: ± SEM between coefficients. **(E)** Average zscored-logged firing rate for novel odors, familiar odors and the difference between novel and familiar stimuli per trials.

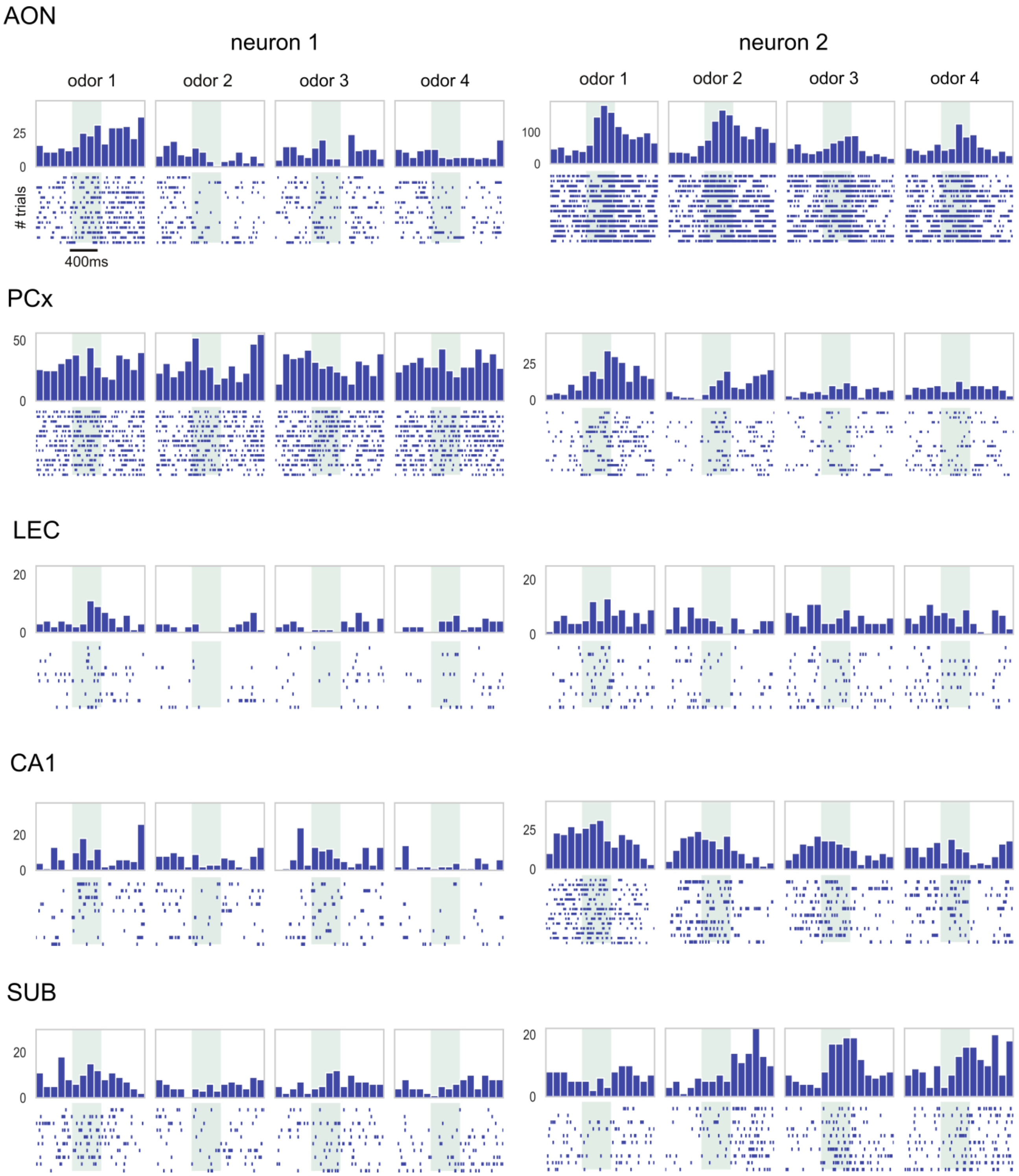


**Figure S3. Example of neurons with high coefficient for the decoding of odor identity.** Example of two neurons per region with high coefficient (>90%).

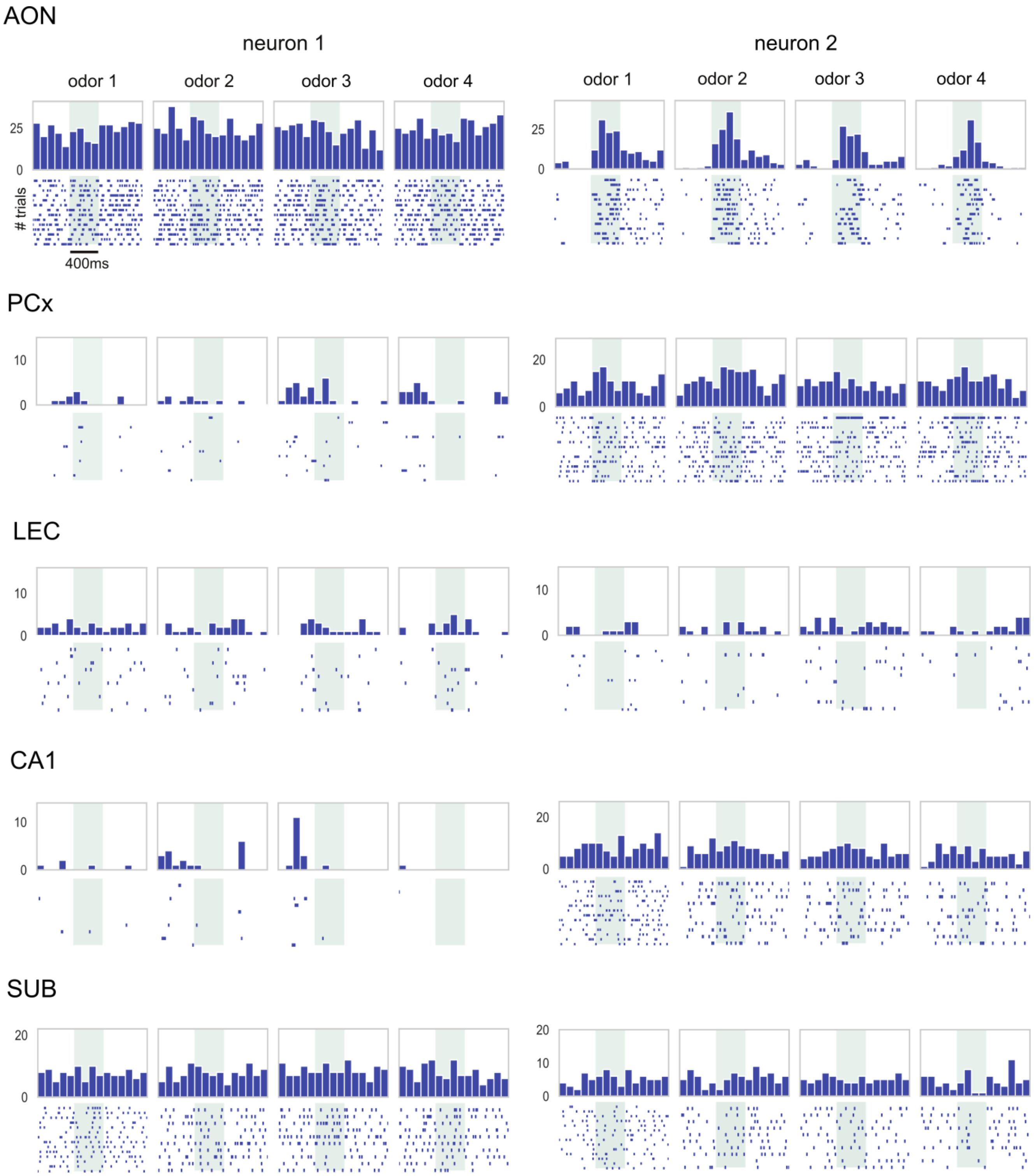


**Figure S4. Example of neurons with low coefficient for the decoding of odor identity.** Example of two neurons per region with low coefficient (<10%).

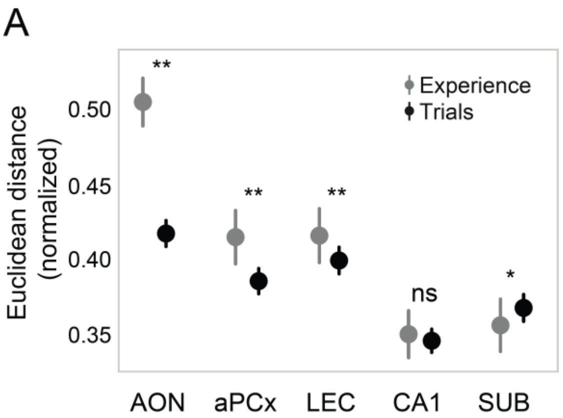


**Figure S5. Quantification of distance in neural space. (A)** Euclidean distance between novel and familiar trials (*Experience*) and between all trials irrespective of experience (*Trials*). In AON, aPCx and LEC, the distance between novel and familiar trials was significantly higher than when not taking experience into account. Conversely, in the SUB, novel and familiar trials were significantly closer than the average distance between all trials (*ns*: not significant, $^{*}p<0.05$, $^{**}p<0.01$). Error bars: Standard error of the mean (SEM).
